# Phenotypic and multi-omics characterization of *Escherichia coli* K-12 adapted to quaternary ammonium compounds identifies lipid A and cell envelope alterations regulated by mar-sox-rob and stress inducible pathways

**DOI:** 10.1101/2020.07.13.201814

**Authors:** Kari A.C. Green, Branden S.J. Gregorchuk, Shelby L. Reimer, Nicola H. Cartwright, Daniel R. Beniac, Shannon L. Hiebert, Timothy F. Booth, Carmine J. Slipski, Patrick M. Chong, Garrett R. Westmacott, George G. Zhanel, Denice C. Bay

## Abstract

Quaternary ammonium compounds (QACs) benzalkonium (BZK) and cetrimide (CET) are common disinfectants used to inhibit or eradicate Gram-negative bacteria in clinical and agricultural products. QAC tolerance in *Escherichia coli* and other Enterobacterales species can confer cross-resistance to various clinically used antibiotics, making it important to understand mechanisms of QAC tolerance in greater depth. QAC adaptation by *E. coli* is hypothesized to alter MarRAB regulated genes that converge on the outer membrane, specifically, lipid A biosynthesis and transport genes, porins, and efflux pump systems. To test this, we performed a ‘multi’-omics and phenotypic characterization of *E. coli* K-12 adapted to BZK and CET, to assess how QACs alter cell growth, genomics, and proteomics. *E. coli* adapted to either BZK and CET resulted in strains with stable QAC tolerance when either drug was omitted, elongated and narrower cell morphologies by scanning electron microscopy, and reduced growth fitness when compared to un-adapted *E. coli*. Antimicrobial susceptibility testing revealed that QAC adaptation increased *E. coli* tolerance by ≥4-fold to BZK, CET, and other QACs but no antibiotic cross-resistance. Single nucleotide variants identified by whole genome sequencing and differentially accumulated proteins by liquid chromatography-mass spectrometry identified alterations to various QAC-adapted *E. coli* genes and proteins belonging to: lipid A biosynthesis and transport (*lpxLM, msbA, mla*), the *mar-sox-rob* regulatory pathway (*marR, rob*), DNA/protein translation (*gyrA, rpsA, rpoB, rapA*). These alterations validate the hypothesis that *mar-sox-rob* network plays a role in QAC tolerance and identifies additional stress inducible genetic and protein QAC tolerant biomarkers.

**Importance:** Bacterial tolerance mechanisms associated with disinfectant QAC adaptation is hypothesized to overlap with the mar-sox-rob multiple antimicrobial resistance pathway but has not been directly shown. Here, we generate QAC tolerant *E. coli* strains and identify phenotypic changes associated with protein and genetic alterations caused by prolonged QAC exposure. We identified genes that overlap with known antibiotic resistance mechanisms as well as distinct genes and proteins specific to QAC adaptation that are useful for future bacterial disinfectant tolerance mechanism studies. However, these altered genes and proteins implicate MarR and Rob pathways specifically in QAC tolerance but, surprisingly, the involvement of mar-sox-rob pathways did not increase antibiotic cross-resistance. Many altered genes we identified were essential genes in lipid A biosynthesis/transport, DNA and RNA transcription, and protein regulation systems potentially explaining why only QAC cross-tolerance was observed and why we observed greater cell fitness costs despite MarR and Rob pathway involvement.

## 1.0 Introduction

Quaternary ammonium compounds (QACs) encompass a wide range of antiseptics, disinfectants, and surfactants that possess one or more permanently positively charged nitrogen molecules with hydrophobic aryl and/or acyl chain moieties. The most commonly used antiseptic QACs are detergent-like compounds including benzalkonium chloride (BZK), and cetrimide (CET) (1). QACs have multiple mechanisms of action to inhibit growth and/or kill bacterial cells and their most well recognized is their ability to disrupt membranes resulting in ion leakage (2). The permanent positive charge(s) of QACs increase reactive oxygen nitrogen species formation and denaturation of proteins, contributing to cell death (1, 2). QACs are widely used in clinics (3), food preparation facilities (4), households (5), agriculture/aquaculture (6), and industrial facilities (7, 8) due to their antimicrobial activity and low cytotoxicity (9). The global over-usage of QACs and their resulting environmental biomagnification have been linked to the emergence and spread of QAC tolerance among clinically relevant antimicrobial resistant Enterobacterales species (10–13). Therefore, it is important to understand how QAC tolerance and antimicrobial resistance mechanisms are linked.

QACs and other antiseptics lack clearly defined resistance breakpoint values based on current Clinical Laboratory Standards Institute (CLSI) guidelines (14), therefore, we refer to QAC resistance as ‘tolerance’ herein. Increased QAC tolerance of Enterobacterales is concerning, as it is frequently associated with and may increase cross-resistance to clinically used antibiotics (15, 21, 22). This is particularly important for *Escherichia coli*, the most common Gram-negative organism identified from blood and urinary tract infections in North American hospitals (15–17) and a frequent food-borne infectious agent in meat processing (18, 19) and water treatment plants (20, 21).

When compared to antibiotic resistance, QAC tolerance mechanisms are not as well-defined in Enterobacterial species (22, 23). Established mechanisms of QAC tolerance converge on cell envelope alterations including: i) upregulation or acquisition of QAC-selective efflux pumps (intrinsic *acrAB* and *mdfA* systems (24) or acquisition of *qac* genes (23–25)), ii) altered expression/accumulation of general diffusion outer membrane porins (*ompC* and *ompF* (26)), and iii) lipopolysaccharide (LPS) and phospholipid modifications (*lpxM* and *lpxL* (2, 27)). In antibiotic resistant species, the expression of these porins (28), efflux pumps (29, 30) and lipid A biosynthesis/transport (31) genes are regulated by global stress response transcription factors that recognize the *mar-sox-rob* box regulon (32) but it remains unclear how QACs influence this pathway. Insight into QAC tolerance mechanisms are most frequently obtained from studies involving laboratory-adapted species (11, 13, 15, 31, 38–40), where only a few studies applied an ‘-omics’ based approach to identifying QAC induced cell alterations. Transcriptomic (33) and proteomic (10, 11, 33, 34) alterations of QAC adapted strains have been reported but no insights into potential genomic alterations are available.

Here, we address the hypothesis that increased QAC tolerance alters Enterobacterial genomes adapted to QACs by altering lipid A, efflux pump and porin, systems controlled by the multiple antibiotic resistance (Mar; *marRAB*) transcriptional regulatory pathway. The MarRAB pathway has been proposed to participate in antiseptic activation in previous studies (4, 35) but was never directly shown. To accomplish this, we gradually adapted the *E. coli* K-12 strain BW25113 to two of the most commonly used QACs, CET and BZK We applied a combination of growth-based assays, imaging, and multi ‘omics’-based approaches to identify the phenotypes, genes, and proteins associated with *E. coli* QAC tolerance. We propose that QAC adaptation in *E. coli* targets and alters specific pathways that converge on the outer membrane, specifically, lipid biosynthesis and transport genes, porins, and efflux pump systems, all of which overlap with known *mar* regulated antimicrobial resistant mechanisms. Hence, QAC-adapted *E. coli* that have a stable QAC tolerance phenotype should be expected to have genetic changes in similar systems or the same genes based on monitoring whole genome sequenced (WGS) single nucleotide variants (SNVs) and altered proteomic protein abundance profiles by whole-cell liquid chromatography mass spectrometry (LC-MS/MS). Growth curves, antimicrobial susceptibility testing (AST), cell imaging by scanning electron microscopy, and QAC-tolerant phenotype stability AST of each adapted *E. coli* were measured to compare cell fitness and morphology to unadapted *E. coli*. This analysis identifies genes and proteins specifically associated with QAC adaptation that may serve as future QAC tolerance biomarkers. It also reveals that *mar-sox-rob* pathway participates in QAC tolerance but is not necessarily associated with antibiotic cross-resistance.

## 2.0 Results

### 2.1 Adapted E. coli replicates showed stable QAC tolerance phenotypes to BZK and CET based on AST

To identify the genetic and proteomic alterations caused by QAC adaptation, we generated three independently adapted replicates of *E. coli* K-12 BW25113 to either BZK (BZKR1-3) or CET (CETR1-3) QACs using daily sub-culturing method that involved stepwise, increasing QAC concentrations over 40 days (Table 1). The AST results of each ‘day 40’ QAC-adapted replicate confirmed that both sets of CETR and BZKR replicates exhibited a 4-to 8-fold increase in their respective QAC MIC values when compared to the unadapted *E. coli* (WT) strain (Table 1). Based on their successful adaptation to BZK and CET, we measured the phenotypic stability of each replicate’s QAC tolerance phenotype. Previous QAC-adapted bacterial experiments have reported that when the QAC selective pressures are removed in growth experiments QAC tolerance phenotypes can be quickly lost (36, 37). To measure QAC phenotype stability, we sub-cultured QAC-adapted replicates without QAC selection over 10 days and performed AST measurements each day (Table 2). The results of this analysis show QAC-adapted replicates maintained a QAC tolerant phenotype until day 10, where they demonstrated a decrease in their respective MIC values. Despite the decreased MIC at day 10, all BZKR and CETR MIC values remained significantly higher (>2-fold) than the WT MIC value (Table 2), indicating that after 40 days, QAC adaptation of *E. coli* K-12 resulted in a relatively stable QAC-tolerant phenotype.

**Table 1.**
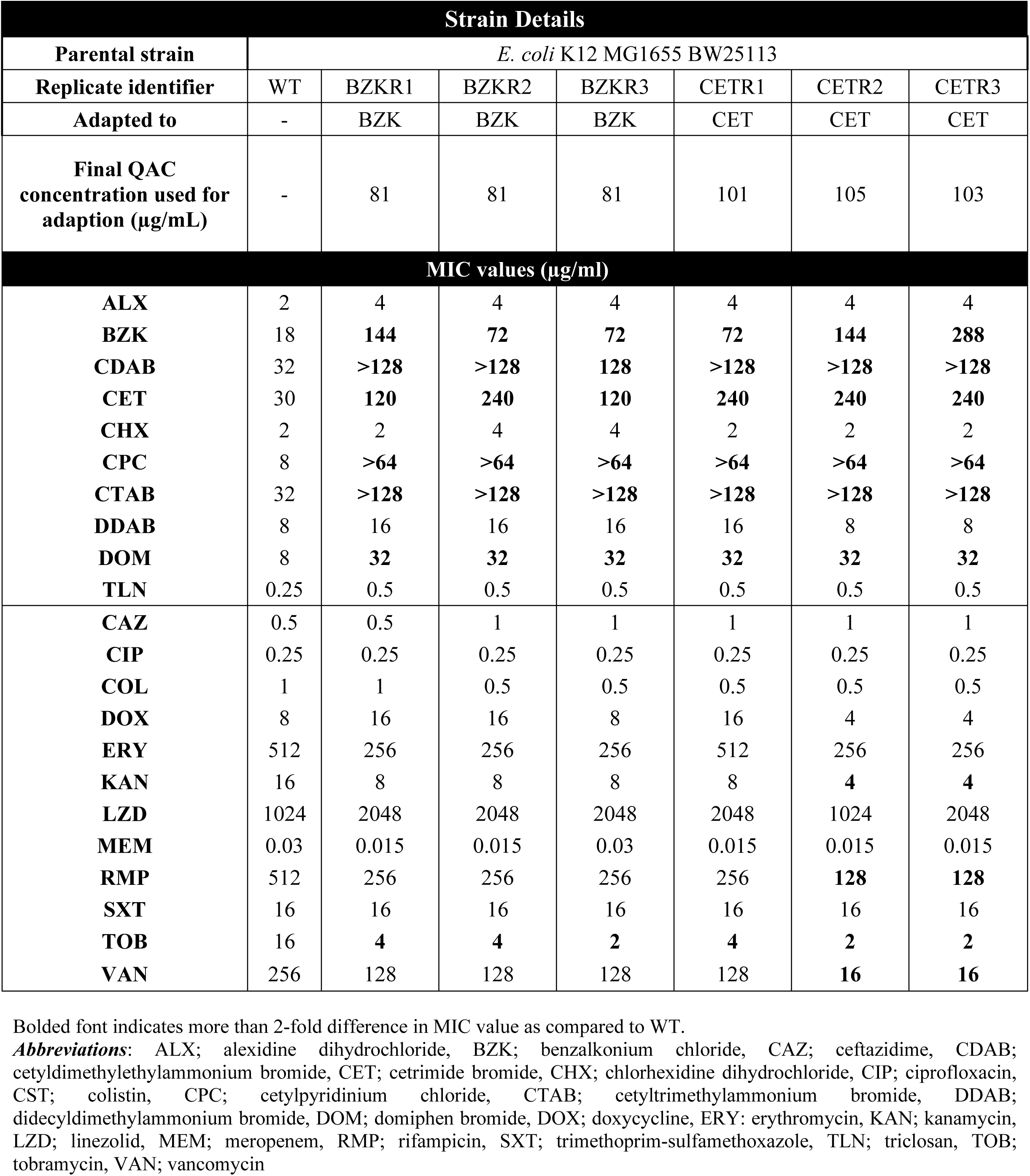
Strain details and mean MIC values for QAC-adapted replicates after 40 days of QAC exposure.

**Table 2.**
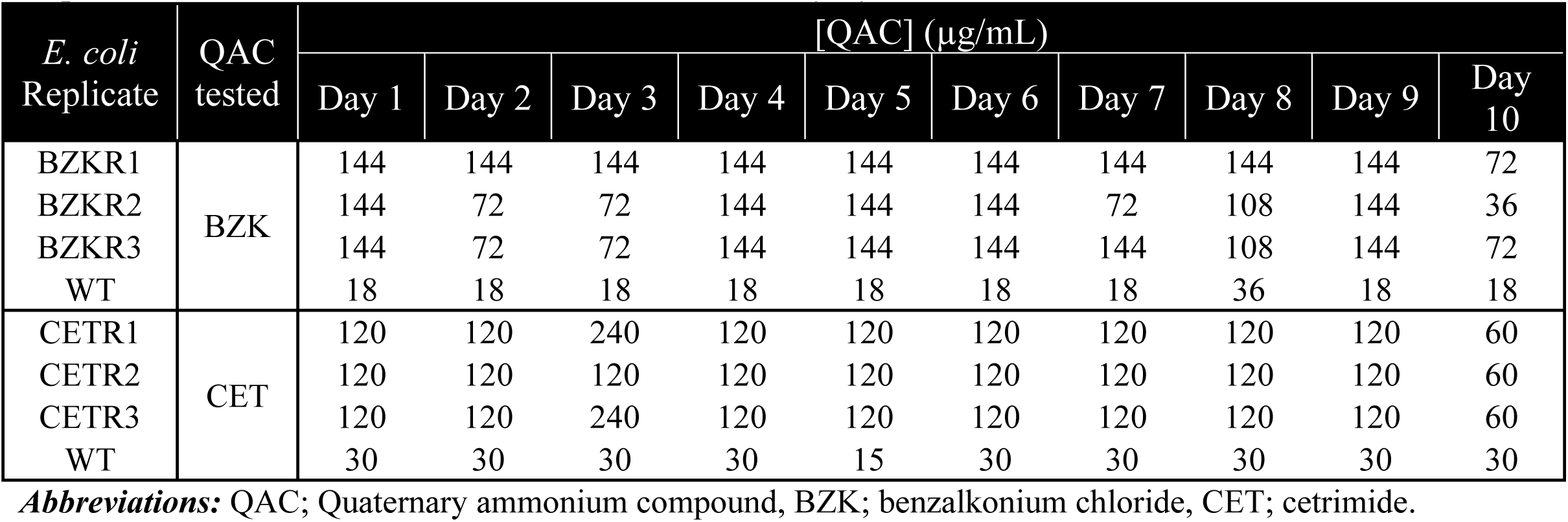
A summary of the stability of the QAC tolerant phenotype in LB broth over 10 days and their respective QAC mean MIC values determined each day by AST.

### 2.2 QAC-adapted E. coli replicates have reduced growth fitness in all media but no significant differences in biofilm growth

Prolonged QAC exposure was previously shown to come at considerable growth fitness costs many adapted Gram-negative species, due to the accumulation of tolerance conferring mutations that can reduce the growth rate of the culture (38–40). To investigate the fitness of QAC-adapted *E. coli*, 24 hr growth curve experiments were performed for all replicates in various rich and minimal media with and without added sub-inhibitory QAC concentrations (Fig. 1, Fig. S1-S2). After comparing growth curves in all media, the results indicated that BZKR replicates were approximately 40% slower in growth rate as compared to WT and CETR replicates were approximately 30% slower than WT (Fig. 1, S1-S2). Both QAC-adapted *E. coli* replicates grown in Mueller Hinton broth (MHB) at sub-inhibitory QAC concentrations respective to the WT attained a 25-30% reduction in final OD_600nm_ value after 16 hrs when compared to the WT or the respective replicates grown in MHB only (Fig. S1). Both BZKR and CETR replicates grown in any media with drug (except for minimal nine salts; M9) had 1.3-2.2 fold reduced doubling time when compared to the same replicate grown in the same media without drug (Fig. S1-S2). This indicated that QAC addition to nearly all media did not confer any specific growth advantage to the adapted replicates. All QAC-adapted replicates grown in minimal media (+/-QAC) showed a 30-40% decrease in final optical density at 600 nm (OD_600nm_) values after 24 hr growth (maxima at 0.5-0.9 units) and exhibited a longer lag phase (2-4 hrs) when compared to WT grown under the same conditions (Fig. 1, Fig. S2). Together, this indicates that the fitness of QAC-adapted *E. coli* was measurably reduced in nearly all media when compared to WT, particularly when grown in minimal nutrient defined media.

**Figure 1.**
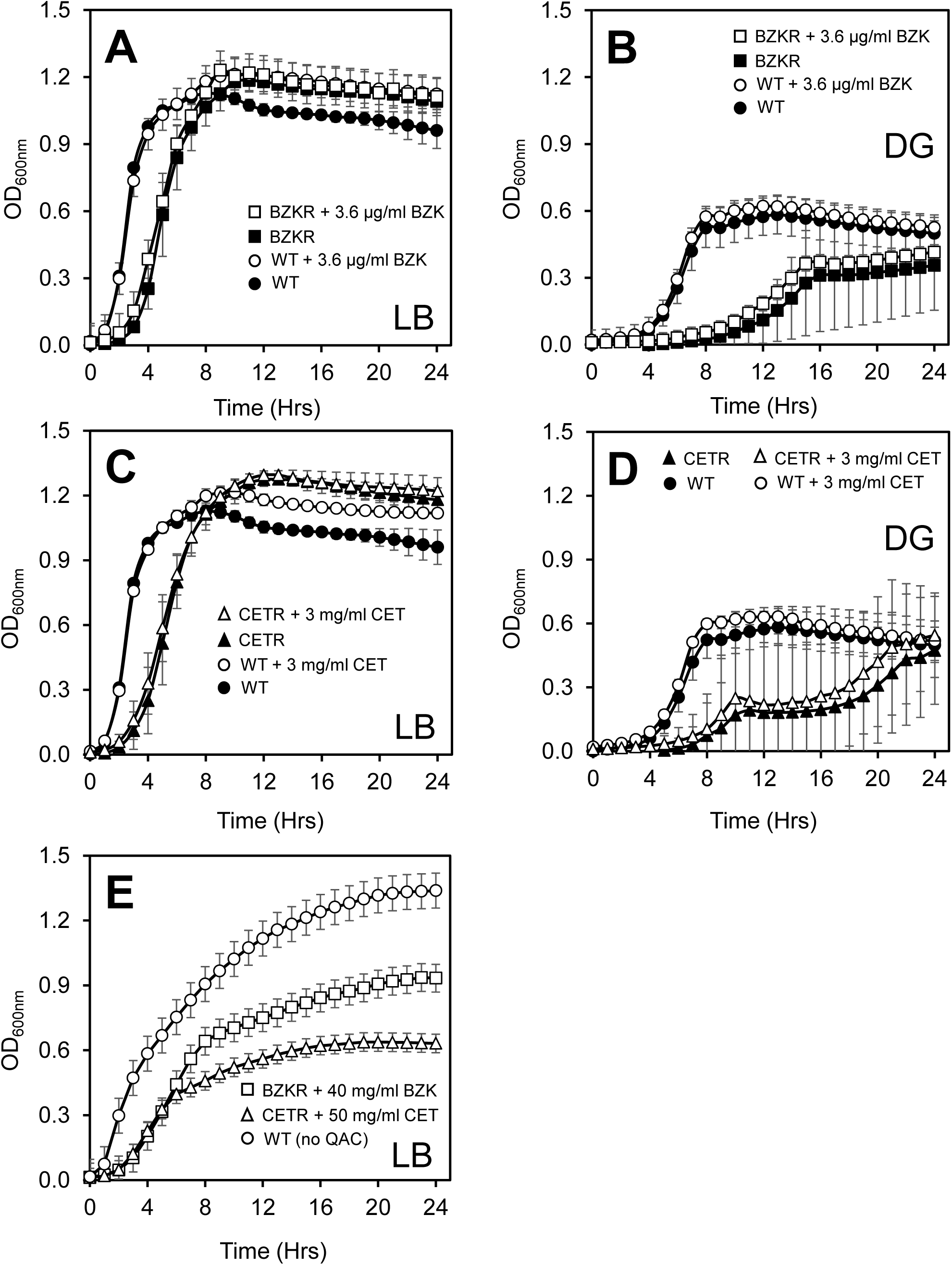
24 hr growth curves of QAC-adapted *E. coli* grown in LB or DG media in the presence and absence of sub-inhibitory QAC concentrations. Panels **A-B** show growth curves of BZK-adapted replicates (squares) and WT (circles) grown with (white) or without (black) 3.6 µg/ml BZK in LB (**A**) or DG (**B**) media. Panels **C-D** show growth curves of CET-adapted replicates (triangles) and WT (circles) grown with (white) or without (black) 3 µg/ml CET in LB (**C**) or DG **(D**) media. **E**) 24-hour growth curves of BZK-adapted (open squares), and CET-adapted (open triangles) replicates with selection (40 µg/ml BZK; 50 µg/ml CET) against WT (open circles) in LB media only. In panels **A-D**, each graph shows the mean OD_600nm_ of the three adapted biological replicates (n=3), in panel **E** each replicate was inoculated in technical triplicate (n=9) and the error bars shown for each measurement represent the standard deviation at each time point.

Based a on previous study, biofilm formation can increase the QAC tolerance of bacteria (41), thus, we investigated if CETR or BZKR replicates had enhanced biofilm biomass formation (Fig. S3). Crystal violet biofilm formation assays of *E. coli* biofilms grown on the pegged lid of a MBEC device, indicated that there was no significant difference in biofilm biomass formation between the QAC-adapted replicates and WT (Fig. S3). There was a significant (*P*<0.05) difference in the biofilm biomass between the BZKR and CETR replicates themselves. This finding indicates that both QAC-adapted *E. coli* produced similar levels of biofilm formation when compared to WT and reveal that *E. coli* biofilm formation is not significantly altered by QAC adaptation.

### 2.3 BZK and CET adaptation cause morphological changes in cell length and width

To determine if prolonged QAC adaptation resulted in cell morphology changes, QAC-adapted *E. coli* was compared to the WT by scanning electron microscopy (SEM). Each QAC-adapted replicate was visualized at mid-log phase (OD_600nm_ = 0.5) and compared to WT to determine differences in cell length/width measurements from SEM images at 5000X magnification. Representative SEM images of selected QAC-adapted replicates show that these cells had a characteristic bacilliform cell shape but also differences in cell dimensions and surface morphology when compared to the WT (Fig. 2A-C). Cell length and width measurements of each QAC-adapted replicate in Fig. 2 D-E revealed measurable but statistically insignificant increases in cell length for CETR3 (+0.085 µm), BZKR1 (+0.056 µm), and BZKR2 (+0.085 µm) when compared to WT (1.826 µm ± 0.038 µm). Significant differences in length were noted for BZKR3 (+0.376 µm), CETR2 (+0.194 µm), and CETR1 (−0.379 µm) as compared to the WT. All QAC-adapted replicates had significantly decreased cell width (BZKR1-3 strains; -0.143 to -0.287 µm; CETR1-3 strains: -0.142 to -0.179 µm) when compared to WT (0.871 µm ± 0.090 µm). These findings indicate that QAC adaption in *E. coli* resulted in narrower cells that varied in cell length but all cells maintaining a bacilliform shape.

**Figure 2.**
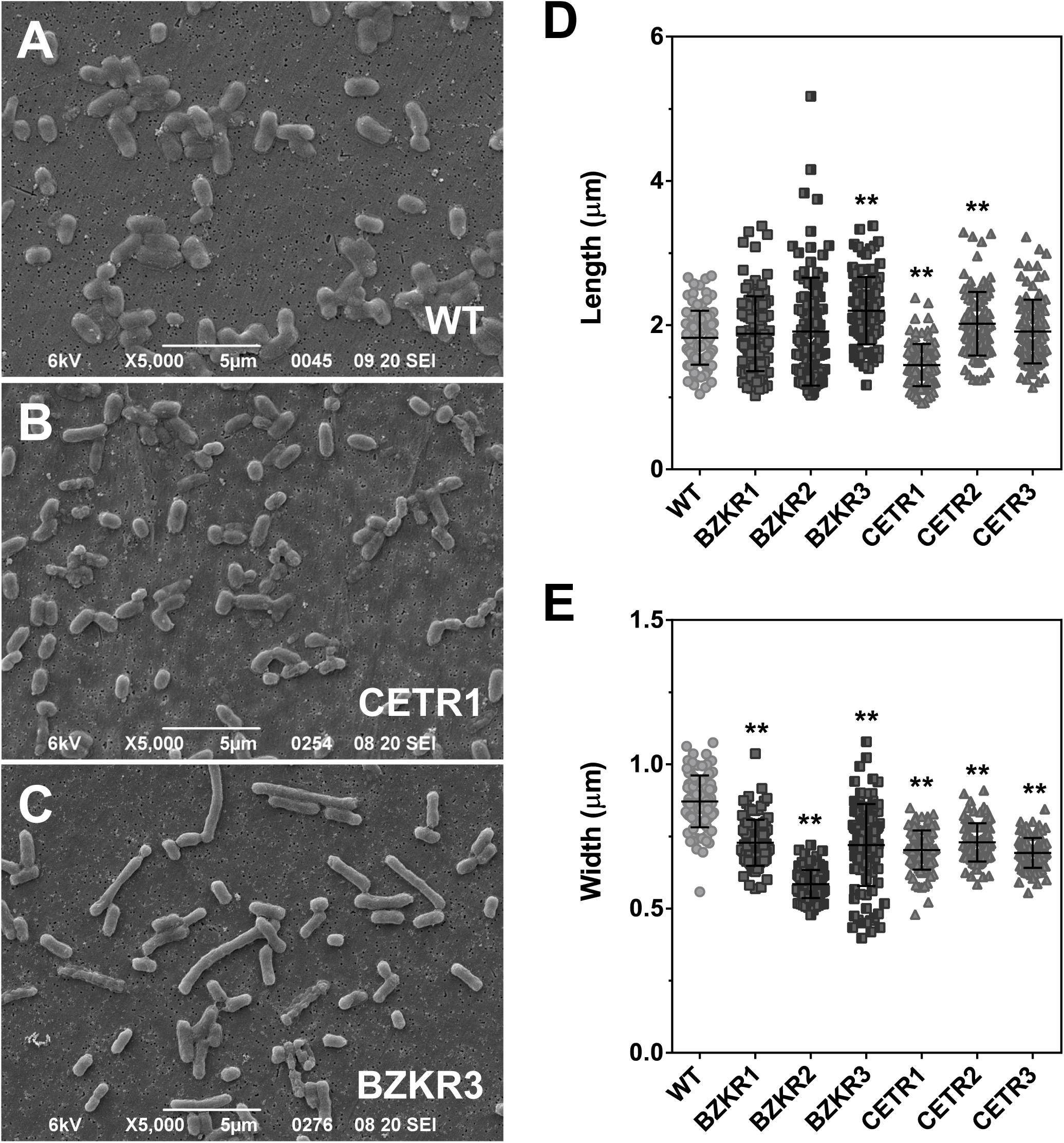
Cell morphology, width and lengths of mid-log *E. coli* strain SEM images at 5000X magnification. Representative SEM images of WT (**A**), CETR1 (**B**) and BZKR3 (**C**) are shown. Length (**D**) and Width (**E**) measurements for QAC-adapted *E. coli* cells (n=100) are shown separated according to replicate measured from SEM images. In panels D-E, asterisks (**) represent significantly different length/width values between a replicate compared to WT (*P* < 0.01), and error bars represent the standard deviation of the mean measured values.

### 2.4 QAC-adapted E. coli demonstrated cross-tolerance to QACs, where some replicates had greater susceptibility to certain antibiotics

Studies of QAC-adapted bacteria have previously shown that QAC adaptation increases tolerance/resistance not only to the compound used for adaptation, but also to additional antimicrobials (10, 11, 40). To evaluate antiseptic cross-tolerance/antibiotic-resistance, AST was performed with a collection of antimicrobial agents listed in Table 1. QAC-adapted replicates had significant cross-tolerance toward other QACs (>2-fold increased MIC), with the exception of DDAB when compared to the WT. Significant cross-tolerance to other common antiseptics, such as chlorhexidine (CHX), alexidine (ALX), and anionic biocide triclosan (TLN) was not observed by any QAC-adapted replicate (Table 1), highlighting the specificity of QAC adaptation and cross-tolerance. Increased antibiotic cross-resistance was not observed by either QAC-adapted *E. coli* (Table 1). However, all QAC-adapted replicates showed a significant (4-to 8-fold) increase in susceptibility to tobramycin (TOB), while CETR2 and CETR3 demonstrated increased susceptibility to Gram-positive selective antibiotics, rifampicin (RMP) and vancomycin (VAN). Together, our results indicate that adaptation of *E. coli* to BZK or CET significantly enhances cross-tolerance to all but one QAC we examined. Additionally, QAC adaptation did not provide significant (>2-fold) cross-resistance to any antibiotics tested. Hence, the QAC-adapted *E. coli* replicates confer enhanced tolerance to other QACs, making these *E. coli* replicates ideal for studying QAC tolerant mechanisms of tolerance.

### 2.5 QAC-adapted E. coli possess repetitive SNVs in genes associated with lipid A biosynthesis, transport systems, and marR and rob stress induced transcriptional regulators

To determine the extent of genetic alterations that occurred in each QAC-adapted replicate, we sequenced each replicate’s genome using WGS to identify SNV differences from the WT genome. SNVs in coding (gene) and non-coding (intergenic) regions of each QAC-adapted replicate were detected, and their genetic relatedness was compared by Maximum Likelihood phylogenetic analysis (Fig. 3A). Phylogenetic analysis revealed that BZKR1 and BZKR3 as well as CETR2 and CETR3 branched closely together and had similar SNV alterations, whereas CETR1 branched furthest apart. CETR1 had the highest number of coding SNVs in the fewest repeatedly identified genes while BZKR2 had the fewest SNVs (Fig. 3A; Table S2).

**Figure 3.**
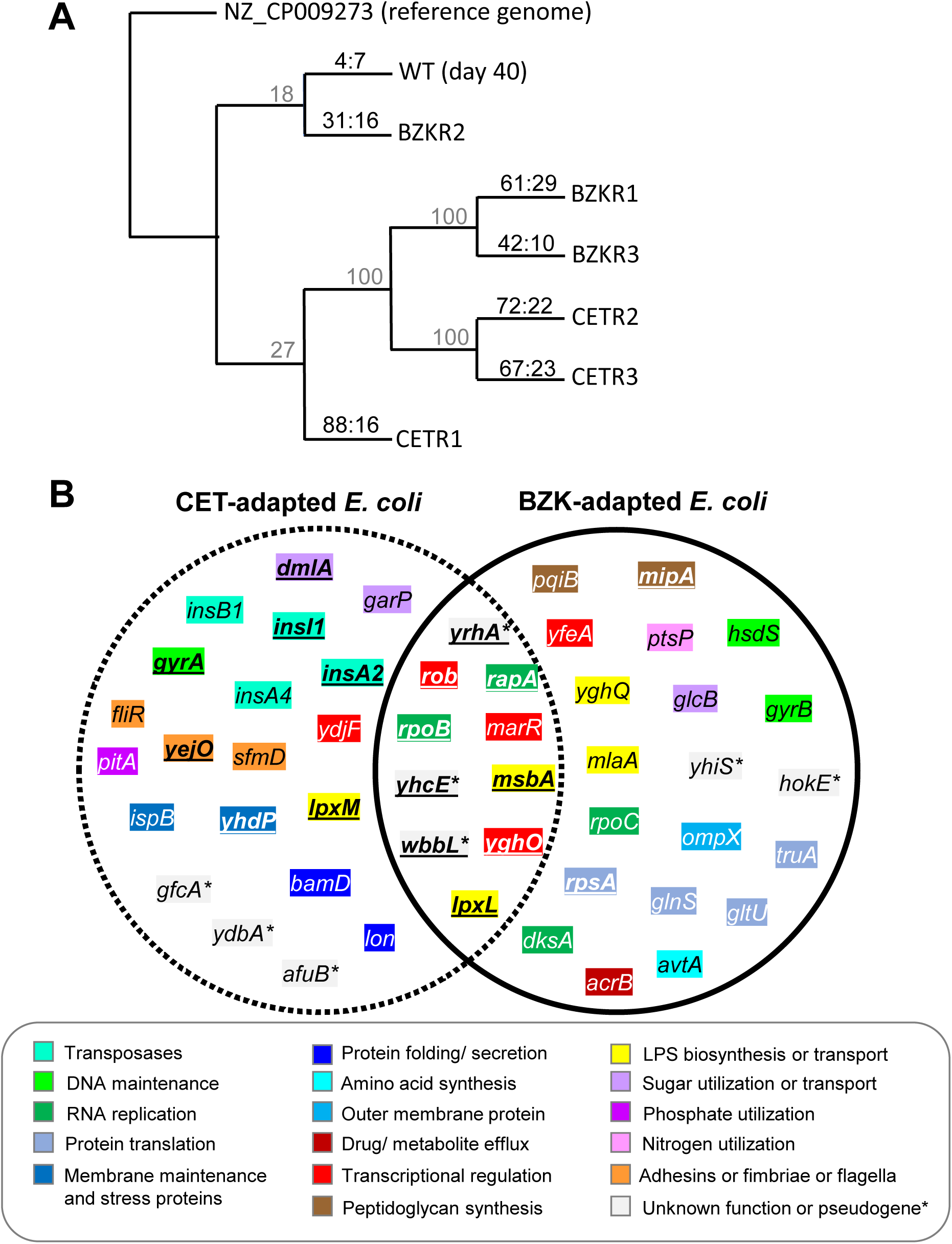
Summary of SNV analyses for each WGS QAC-adapted *E. coli* K-12 replicate. **A)** Maximum likelihood phylogenetic tree for QAC-adapted replicates based on SNVs. Percentage bootstrap (n=1000) support value are shown in grey font above each node. The ratio of coding to non-coding (coding: non-coding) SNVs identified in each genome are listed on each branch according to its respective QAC-adapted replicate. WT (day 40) refers to the WT *E. coli* BW25113 strain cultured without selection for 40 days as a SNV genetic drift control caused by prolonged growth in LB medium only. **B)** A Venn diagram of repeatedly identified SNVs found in coding regions among *E. coli* strains adapted to BZK and CET. Bolded and underlined genes indicate genes with SNVs identified in two or more independently QAC-adapted strains.

In an effort to identify genes that may represent QAC biomarkers, we focused on repetitively identified SNVs occurring in similar coding regions (genes); *i.e*. SNVs occuring in 2/3 or 3/3 BZK-or CET-adapted replicate genomes (Fig. 3B, Table 3). For CET-adapted *E. coli* genomes, repeatedly detected SNVs occurred in genes corresponding to sugar utilization (*dmlA*), outer membrane permeability (*yhdP*), transposase insertion elements (*insA1, insI1*), adhesins (*yejO*), and lipid A biosynthesis (*lpxM*) (Table 3). BZK-adapted *E. coli* genomes had SNVs in fewer repeatedly detected genes, specifically, genes associated with peptidoglycan synthesis (*mipA*) and protein translation (*rspA*) (Table 3). Comparing all QAC-adapted genomes together in a Venn diagram revealed a total of 8 genes that possessed repeatedly identified with SNVs in at least 2/3 replicates (Fig. 3B): the biofilm associated transcriptional regulator (*yghO*), RNA polymerase genes (*rpoB, rapA*), lipid A biosynthesis enzyme (*lpxL*), and lipid A ATP-dependent flippase transporter component (*msbA*). Three pseudogenes had repetitive SNVs identified in the QAC-adapted genomes, corresponding to LPS O-antigen modification (*wbbL1-2*), chaperone-usher fimbriae (*yhcE*), and a pseudogene of unknown function (*yrhA1-2*) (Fig. 3B, Table 3). SNVs occurring in genes belonging to the same operon were also observed among the QAC-adapted replicates. For example, the DNA gyrase operon *gyrAB* and the RNA polymerase β’ subunit operon *rpoBC* had SNVs in both genes among either BZKR or CETR genomes (Fig. 3B, Table 3, S2). These findings indicate that only a few genes with repeatedly identified SNVs were shared by both BZKR and CETR replicates. However, among those SNVs that were shared by multiple replicates, SNVs frequently occurred in lipid A and outer membrane protein associated genes reflecting alterations to genes affected by membrane disruptive QAC mechanism of action. It is important to note that non-deleterious, non-synonymous SNVs were detected in genes identified from previous transcriptomic and proteomic studies such as porin *ompX* (33) (in BZKR3 only) and efflux pump system component *acrB* in BZKR1 (10). Since neither SNV was deleterious and only present in a single QAC-adapted replicate, these SNVs appear to occur less frequently and likely avoid altering these QAC mechanisms of tolerance (Fig. 3B, Table 3).

**Table 3.**
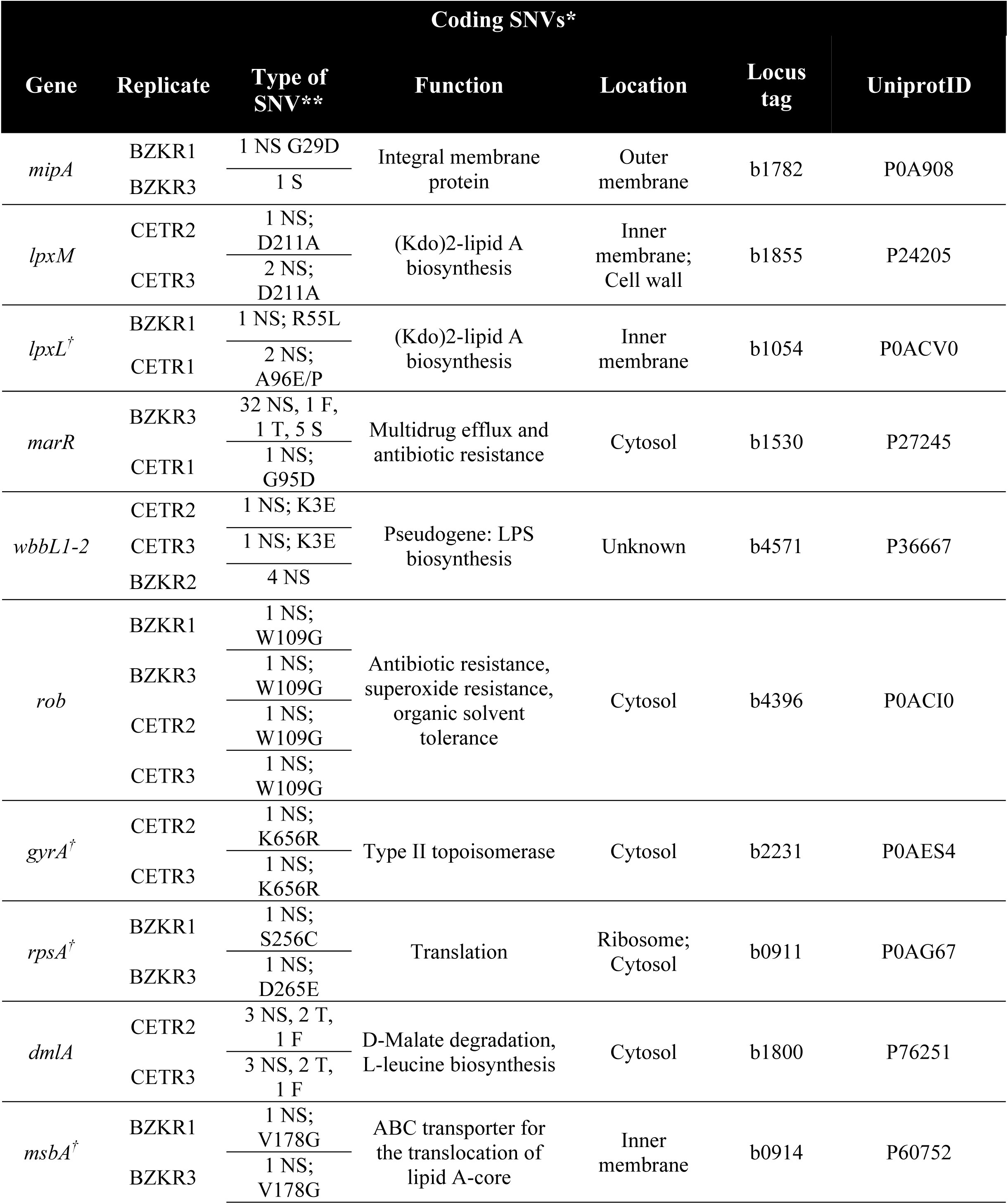
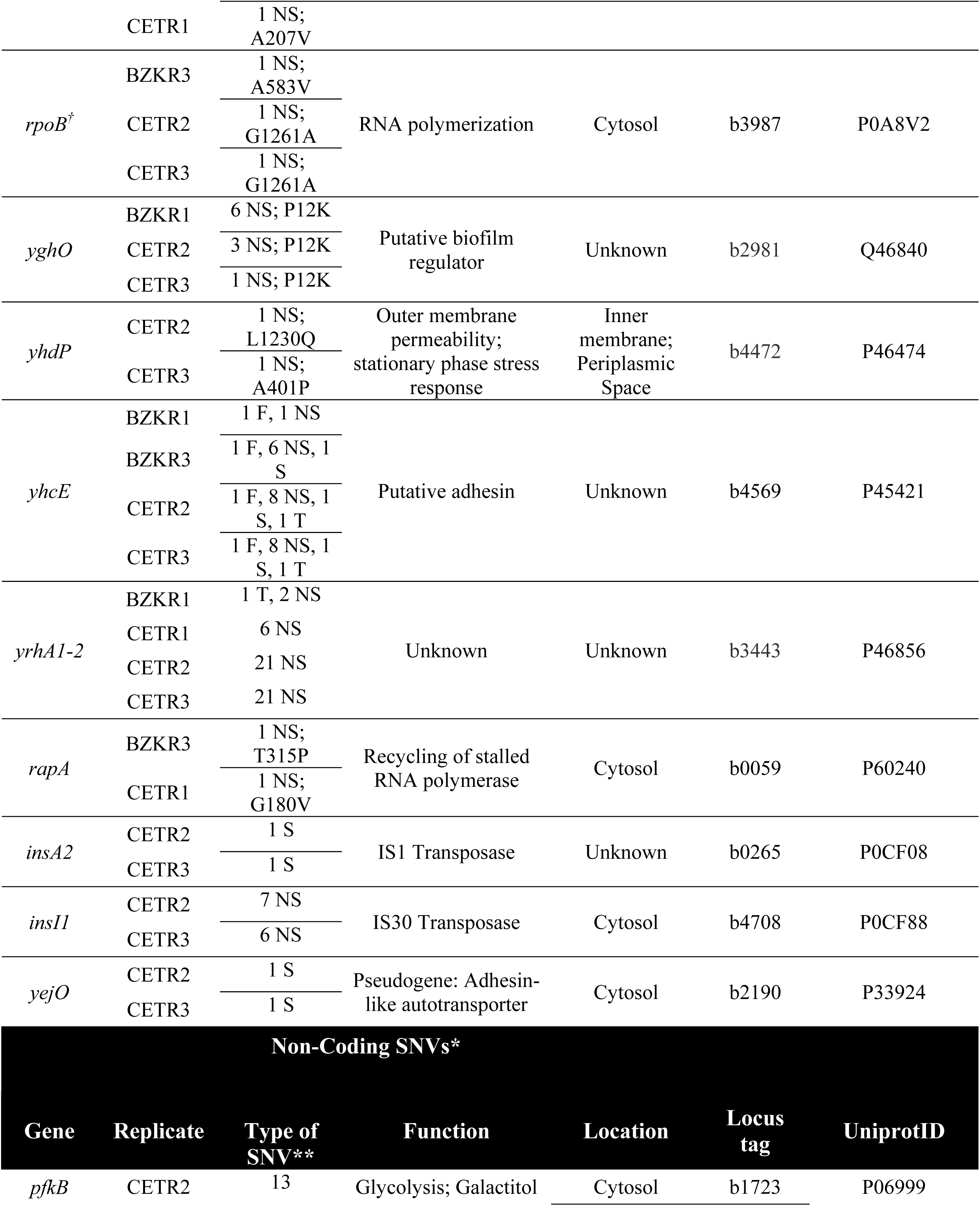
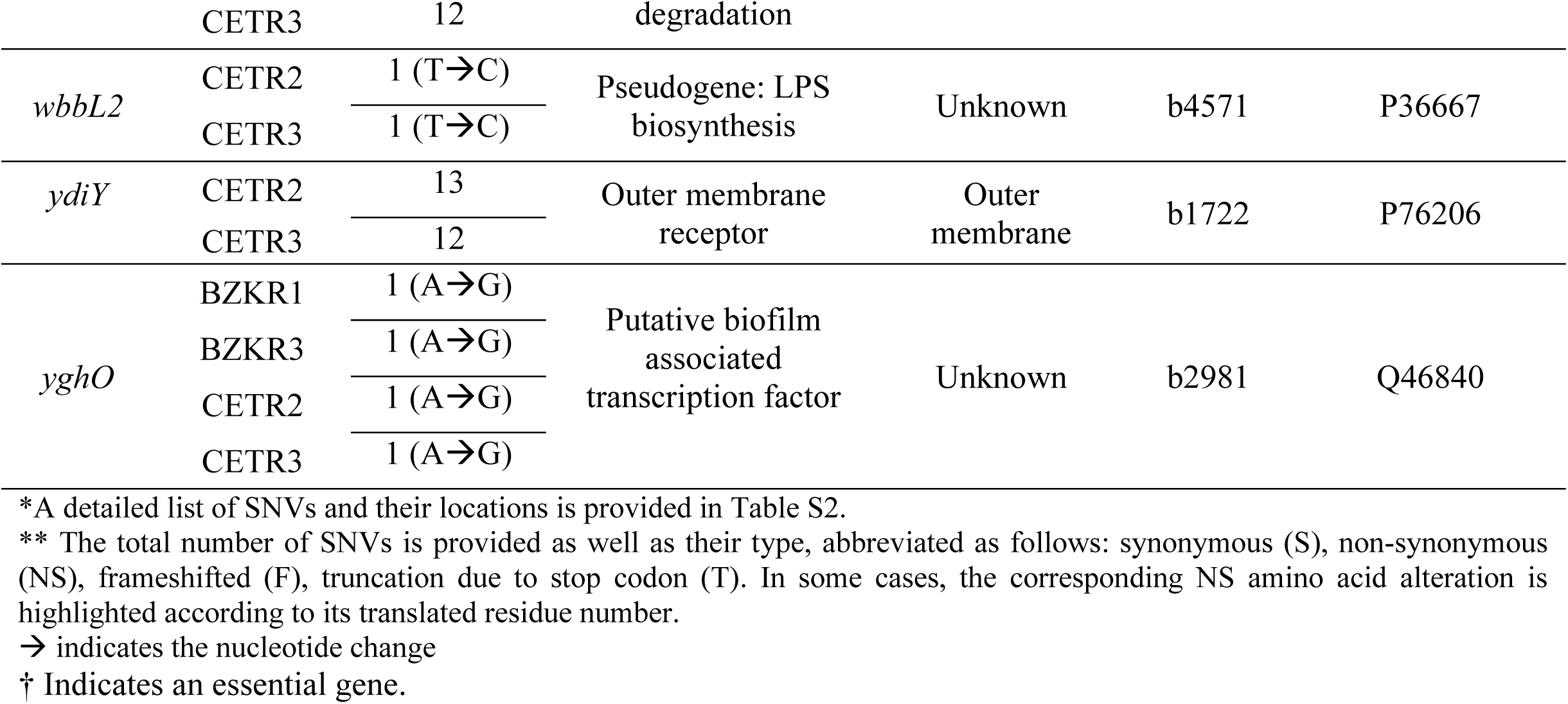
Summary of repeatedly identified SNVs in coding and non-coding regions of each QAC-adapted *E. coli* K-12 genome sequenced in this study.

Lastly and most importantly, we identified deleterious SNVs in *marR*, belonging to the multiple antimicrobial resistance operon *marRAB* operon in CETR1 and BZKR3 adapted strain genomes (Table 3). Additionally, in 4/6 of the QAC-adapted *E.coli* genomes, we detected the same non-synonymous SNV in the transcriptional dual regulator *rob* (Table 3); this SNV generates a mutation that alters Rob amino acid W109G in the predicted turn region after the helix-turn-helix DNA ‘reading’ helix αF based on the crystalized Rob protein structure (42). Identified SNVs in both MarR and Rob indicate that QAC adaptation specifically targets *mar* and *rob* genes directly, validating the hypothesis. SNV alterations to *marR* and *rob* genes would be expected to alter the expression of genes controlled by transcriptional *mar-sox-rob* box regulons, supporting the hypothesis.

### 2.6 QAC-adapted E. coli possesses SNVs in non-coding regions associated with biofilm induced yghO transcriptional regulator and outer membrane genes

Fewer SNVs were detected in non-coding regions of BZKR and CETR replicate genomes than coding SNVs within putative regulatory regions located upstream of a gene (Fig. 3A, Table 3). Only one repeatedly occurring non-coding SNV was detected in 2/3 QAC-adapted replicates and was located within the ribosome binding site region of the biofilm associated transcriptional regulator *yghO* (Table 3, S2). The remaining repeatedly identified non-coding SNVs were identified only in CETR replicates (CETR2-3) and had SNVs in the upstream region of the glycolysis gene *pfkB*, the acid-inducible outer membrane protein *ydiY*, and the insertion element interrupted LPS biosynthesis protein *wbbL1-2* (Table 3). Overall, many of these SNVs were identical in the QAC-adapted replicates, seemingly highlighting their importance in QAC tolerance. It is noteworthy that individual BZKR and CETR replicates had non-coding SNVs in the upstream regions of genes that we identified to have coding SNVs such as porin *ompX* (BZKR3), insertion element *insA* (CETR1), *lon* protease (BZKR1), and retrograde LPS transport system component *mlaA* (CETR1) (Table S2). These SNVs identify genes and non-coding regions that may be useful biomarkers of QAC adaptation and reveal that most relate to outer membrane function or have *mar-sox-rob* regulatory association.

### 2.7 Proteomic analysis identifies proteins with significantly altered accumulation of porins, efflux pumps, and lipid transporter systems associated with mar-sox-rob regulons and other stress inducible systems

Based on the outcome of our SNV analyses, we wanted to verify how the SNV genetic alterations corresponded to potential protein accumulation changes in QAC-adapted *E. coli*. To accomplish this, we performed in-depth whole-cell LC-MS/MS proteomic analysis of CETR1 and BZKR1 replicates as compared to the WT strain. Out of a total of 1904 detected and annotatable proteins from this technique, only 283 proteins in BZKR1 and 161 proteins in CETR1 had significantly altered protein abundance as compared to WT (Table S3; Fig. 4). Only Rob was significantly increased in BZKR1 proteome, whereas MarRAB or SoxSR could not be confidently detected or were absent in the QAC-adapted proteomes. For BZKR1 proteomes, only 4 proteins, Rob, LpxL, AcrB, MsbA, had a SNV in their coding region and 2 proteins (Lon, Rob) had a SNV in their non-coding region (Table S3.1). For CETR1, there were no SNVs in coding regions for significantly detected proteins and only 1 non-coding SNV that had altered protein expression (MlaA) (Table S3.2). This suggests that SNVs appeared to correspond to altered protein accumulation in each QAC adapted replicate.

**Figure 4.**
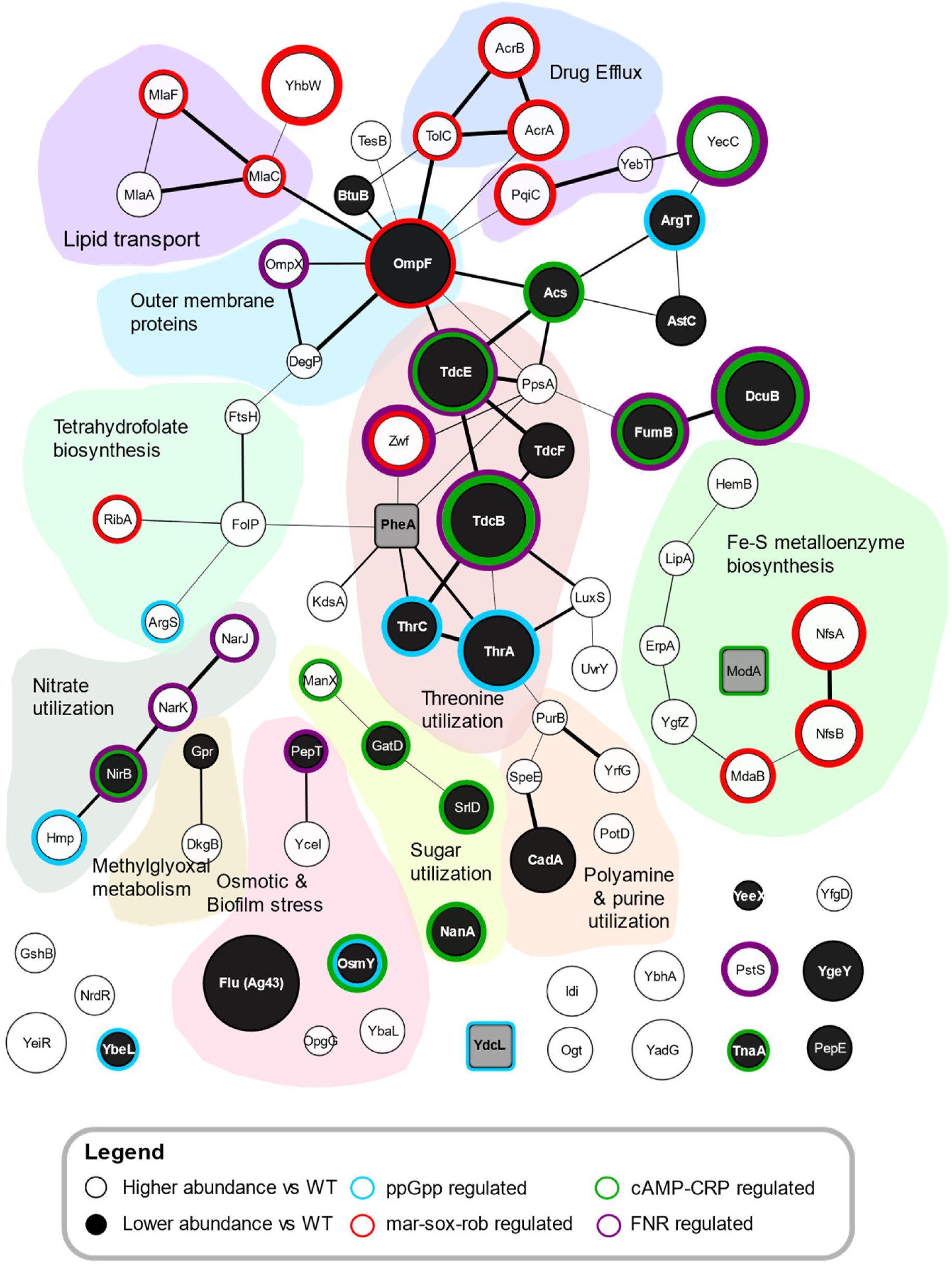
Whole cell proteome STRING protein-protein network diagram of the significantly altered proteins shared in common by BZKR1 and CETR1. Circle nodes indicate each protein and are coloured according to the legend. Protein abundance that was found to be higher (white) or lower (black) in both QAC-adapted replicates by comparison to the WT. Node size is proportional to mean fold change difference in protein abundance. Gray squares represent significant protein abundance changes from WT that were opposite in BZKR1 and CETR1 strains. The circle diameter is proportional to its overall STRING score, which is determined by how frequently proteins are mentioned together in publications. The network diagram was generated using the StringApp v1.5.0 in Cytoscape v3.7.2. All values used in this analysis are summarized in Table S3.

Among all significantly altered proteins, a total of 79 proteins were shared by both CETR1 and BZKR1, with only three proteins (PheA, ModA, YdcL) that were discordantly accumulated between each replicate proteome (Fig. 4). Based on these shared proteins, we identified increased protein accumulations of the efflux pump complex AcrAB-TolC, differential expression of outer membrane porins OmpF (down-regulated) and OmpX (up-regulated), and increased lipid transport system components (MlaACF, PqiC, YebT) in both CETR1 and BZKR1 proteomes (Fig. 4). Among these 79 shared proteins, 13 were *mar-sox-rob* regulated and identified proteins not only involved outer membrane biogenesis and efflux mechanisms, but also those involved in tetrahydrofolate and iron sulfur metalloenzyme biosynthesis pathways (Fig. 4). Differentially accumulated proteins associated with other global stress induced regulatory systems were also noted in Figure 4. Specifically, the starvation and stress inducible stringent response molecule guanosine 3’-diphosphate 5’-diphosphate (ppGpp) (43), the oxygen starvation regulator FNR (44, 45), and finally the secondary carbon source regulator cyclic AMP (cAMP) catabolite gene activator protein (CAP) (46, 47). Hence, *E. coli* adaptation to CET and BZK alters protein accumulation of not only *mar-sox-rob* regulated genes as predicted in our hypothesis, but also additional proteins regulated by other stress inducible protein networks.

To identify changes in functional pathways among the remaining significantly altered proteins of each QAC-adapted proteome, analysis involving the Kyoto Encyclopedia of Genes and Genomes (KEGG) database was performed (Fig. 5). KEGG pathway analysis of the 283 proteins in BZKR1 and 161 proteins in CETR1 could only functionally annotate 56.8% (BZKR1) and 53.8% (CETR1) into a specific ontology (Fig. 5). KEGG analysis highlighted significantly altered protein pathways associated with numerous amino acids, glycerophospholipids, starch and sugar, nitrotoluene, quorum sensing, nucleobase metabolic pathways (Fig. 5). It is noteworthy that many lipid modifying and transport pathways had significantly altered protein accumulation in each QAC-adapted proteome and includes: fatty acid biosynthesis proteins (FabIAH), glycerophospholipid metabolism enzymes (GlpABC, PlsC, GpsA, GlpQ), lipid transport system components (MlaABCF, PqiBC, YebST), and lipid A biosynthesis enzymes (LpxLM) (Fig 5, Table 4). The altered protein accumulation of so many lipid biosynthesis and transport systems supports the hypothesis. Hence, whole cell proteomic analysis of BZK-and CET-adapted isolates revealed significantly altered protein accumulation of many lipid, amino acid, nucleotide and metalloenzyme biosynthetic pathways that may also explain alterations we observed in cell morphology/ fitness experiments but are also are regulated by various stress inducible pathways.

**Figure 5.**
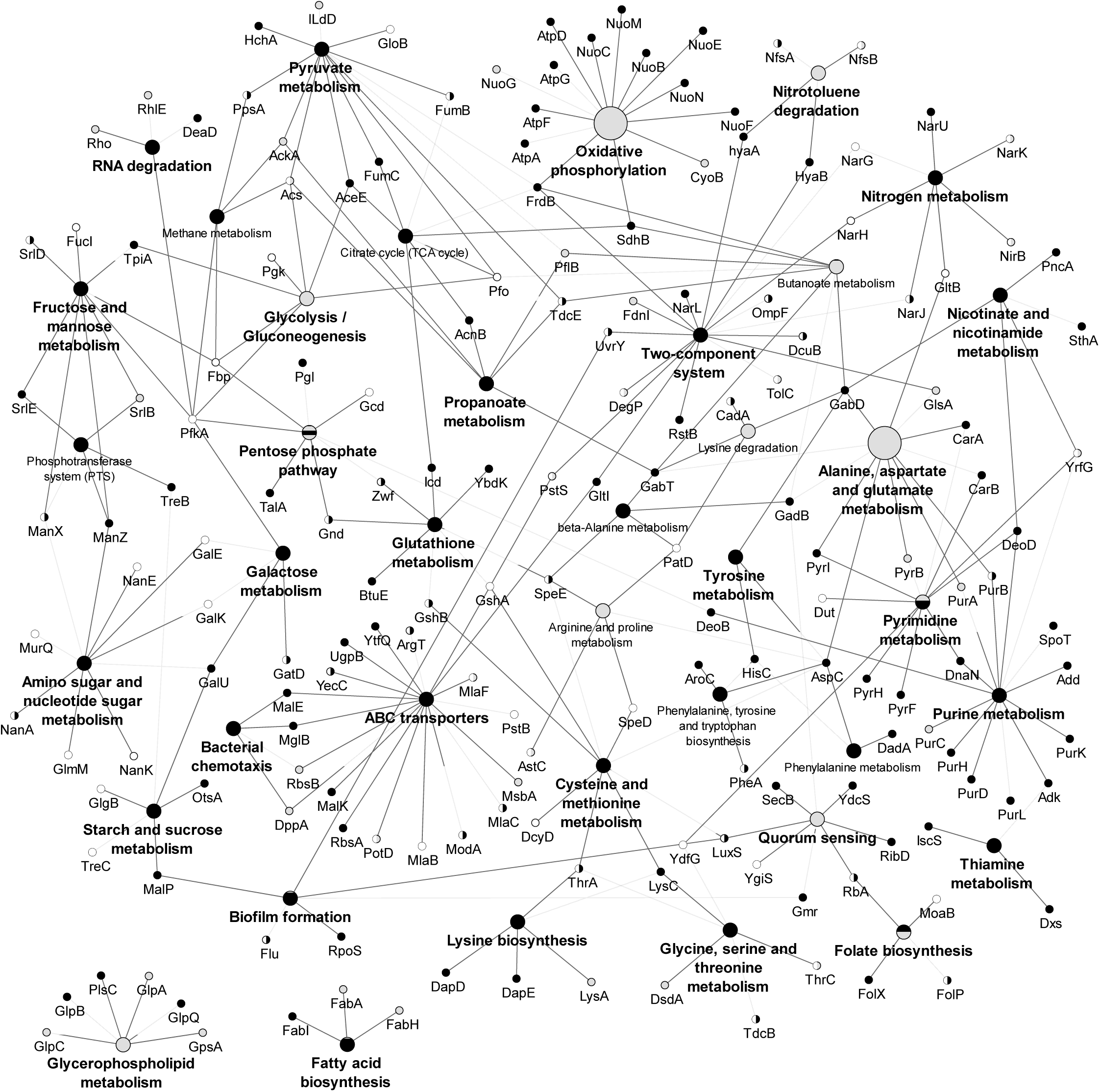
A network diagram summary of whole cell proteomic analysis of BZKR1 and CETR1 with the KEGG database showing significantly altered (up or down regulated) proteins. Significantly altered proteins from BZKR1 (black circles) and CETR1 (white circles) when compared to WT and black/white dots indicate the altered protein was found in both strain proteomes. Lines show connections between the proteins to their common KEGG pathway(s). This network map was generated by Cytoscape v 3.7.2 (96) with the ClueGO v2.5.6 (98) application add-on.

## 3. Discussion

Using a multi-omics approach, this study confirms the involvement of antimicrobial stress inducible transcriptional activators MarR and Rob and the altered accumulation of many *mar-sox-rob* regulated proteins corresponding to efflux pump (AcrAB-TolC), lipid biosynthesis/transport (MlaABCF), and porins (OmpF, OmpX) when *E. coli* K-12 cells are adapted to QACs (Table 2; Fig.3-5). *rob* is part of the *mar-sox-rob* regulon, and is an organic solvent and bile salt inducible transcriptional activator that, along with the *marRAB* operon (48–51), contributes to the overexpression of numerous antibiotic resistance related genes controlled by *mar-sox-rob* boxes, including porin (28) and efflux systems (52). Our study confirms that QAC adaption not only alters *mar-sox-rob* pathway components, but other genes and proteins associated to outer membrane maintenance as well as cellular stress responses, such as lipid A biosynthesis and transport systems (*msbA, lpxLM*), oxidative stress (*yhbW*), DNA maintenance (*gyrAB*), protein translation (*rpsA*), or RNA transcription (*rpoB*C) (Table 3). Alterations to these gene targets highlight the involvement of previously identified intrinsic antimicrobial resistance gene targets, *msbA* (53, 54), *rpoB* (55), *gyrAB* (56, 57), *rpsA* (58); however, only *gyrAB* has been implicated in tolerance to other disinfectants/antiseptics such as triclosan (57). Alterations of these antimicrobial resistance-associated and essential genes also emphasizes the selective pressure exerted by QAC exposure and reflects the contributions of multiple mechanisms of QAC action (*i.e*. membrane disruption, oxidative DNA/protein damage, and protein denaturation (2)). We also saw mutations in many pseudogenes (*wbbL, yhcE, yrhA*) and transposases (*insA, insI, insH*; Table 3), suggesting that alteration of regions controlling genetic exchange are enhanced by QAC adaptation. Furthermore, transposase and insertion element alterations offer additional context to previous studies that monitoring the mobilization of QAC specific resistance genes on mobile genetic elements (25, 59). Repeated identification of SNVs in these essential genes at identical genetic positions in separately adapted *E. coli* replicates (*rob, msbA, rpoB, gyrA, yhgO, ychE*; Table 3; Table S2) revealed the importance of particular codon positions for future antiseptic tolerance biomarker determination. Lastly, less than 1/4 of the genes and proteins detected in this study were previously identified in past transcriptomic and proteomic publications of BZK-adapted *E. coli* (10, 33). Our approach not only identified previously observed QAC targets, but expanded new QAC biomarker detection and confidence by using more than one QAC-adapted replicate.

The results from our various phenotypic assessments of BZKR and CETR growth and QAC tolerance phenotype stability (Fig.1, Tables 2) also appear to agree with previous studies that show QAC adaptation generally comes at a fitness cost to the cell (39, 40, 60). In our growth curve experiments, we observed that QAC-adapted replicates all exhibited delayed growth when compared to WT (Fig. 1, S1-S2). We also observed that, after 10 days of non-selective sub-culturing, the stability of QAC MIC values reduced only after day 9 (Table 2). Previous studies have shown that QAC tolerant phenotypes can be lost when selective pressure is removed (61, 62). However, it is important to note that these studies adapted their bacterial sub-cultures over shorter timeframes (≤20 days) than our study (40 days).

Many of the fitness changes we observed are also likely attributed to SNVs in essential genes in the QAC-adapted replicates (Table 2). The numerous and often identical non-synonymous SNVs we observed in essential genes corresponded to lipid A pathways (*msbA, lpxLM*), protein translation (*rpsA*), RNA transcription (*rpoB*), DNA maintenance (*gyrAB*) (Table 3, Fig. 3). Changes to essential genes often impact cell growth and fitness when altered, but they also acquire additional compensatory mutations in closely related associated genes, which we also observed. For example, *msbA* overexpression reportedly supresses the lethality of *lpxL* mutations (63), and we observed SNVs in both *msbA* and *lpxL* in the same replicates (BZR1, CETR1) (Fig. 3-4, Table 3). Furthermore, we observed altered protein accumulation in a variety of metabolic pathways (eg. sugars, nucleotides, lipids, and amino acids) in BZKR1 and CETR1 (Fig. 5). Alterations to metabolic pathways controlling sugars, amino acids and nucleotides caused by prolonged exposure to BZK or CET would also be expected to negatively impact QAC-adapted *E. coli* growth particularly in nutrient limited minimal versus the rich media used to initially adapt the *E. coli* replicates.

SEM visualization revealed cell morphology differences by QAC-adapted *E. coli*, where CETR cells exhibited chained morphology, and elongated BZKR cells when compared to WT (Fig. 2). Hence, cell division problems appear to be a consequence of prolonged QAC adaptation based on our SEM imaging. We identified numerous outer membrane gene/protein alterations that may impact cell morphology: lipid A biosynthesis (LpxLM), LPS flippase protein (MsbA), and phospholipid transporter systems (MlaACF, PqiBC, YebST) as well as genes (SNVs) and proteins related to outer membrane maintenance (DegP, YhdP; Table 2; Fig. 3-4). All of theses systems may alter the location, structure, and/or the net negative charge of LPS (64–67), ultimately influencing cell morphology. These altered lipid A modifying/ transport systems may increase the production of LPS molecules and permit QAC-exposed cells to replace or modify the LPS that are lost or removed by QAC’s membrane disruptive mechanism of action (2). It is also well established that alterations of plasma membrane phospholipids, specifically phosphatidylglycerol (PG) and cardiolipin (CL), that associate with cell division-localization membrane proteins (ProP, MinD, FtsA, SecA) help determine Gram-negative cell size and shape (68). Although, we did not observe any significant differences in protein accumulation of these cell division-localization protein systems directly, we did observe increased ProQ accumulation in CETR1 proteome (Fig. 4-5; Table S3), which controls ProP expression (69). We also observed protein accumulation changes in fatty acid (FabIAH) and glycerophospholipid enzymes (GlpABC, PlsC, GpsA, GlpQ) and phospholipid transport systems (PqiABC, YebST, Mla) of both QAC adapted proteomes (Fig. 5). All of these lipid biosynthetic enzymes and transporters have been previously noted to impact phospholipid biosynthesis (70–73) and would be expected to alter the content and balance of phospholipids when dysregulated in our QAC-adapted *E. coli*. Disrupting phospholipid distributions have contributed to cells that are narrower in size and width (27, 68), as we observed in our SEM analyses. Further experimental analysis of these QAC-adapted bacteria and their lipids in future studies may reveal additional insights into these morphological changes.

Our AST findings also revealed that QAC-adapted isolates could only confer enhanced tolerance to other QACs but not to other antibiotics (Table 1). AST of CET-adapted replicates showed increased susceptibility to Gram-positive selective RMP and VAN antibiotics that *E. coli* and most Enterobacterales are intrinsically resistant toward (77, 78). Susceptibility to RMP and VAN was previously demonstrated by polymyxin E (colistin) resistant *E. coli* and *Acinetobacter baumannii*; colistin is a membrane-acting antimicrobial known to require LPS modifications as part of its mechanism of resistance (79, 80). In our study, we saw repetitive SNVs and significant protein abundance changes that concentrated on LPS/lipid A transport (MsbA) and biosynthesis (LpxLM) pathways, which are known to have impacts on drug permeability (68, 80, 81). We also saw repetitive SNVs and low or no protein detection of the outer membrane protein YhdP in both QAC-adapted replicates (Table 2, Fig.3B, 4); YhdP is involved in envelope maintenance in association with the cyclic enterobacterial common antigen (ECA) and, when deleted in *E. coli*, Δ*yhdP* cells become more sensitive to vancomycin treatment (82).

Finally, despite our observation that *mar* and *rob* pathway involvement and lipid A alterations were present in QAC-adapted *E. coli* genomes and proteomes, we unexpectedly did not observe enhanced antibiotic cross-resistance. Altered cell morphology of both QAC-adapted *E. coli* likely play a major role, but two additional explanations may also contribute. Firstly, the *mar-sox-rob* regulatory pathway alterations observed were not the sole stress inducible regulatory systems we detected. Our study suggests that other stress inducible regulatory systems, such as the ppGpp regulated stringent response system, the FNR system, and cAMP-CRP system may play a role in regulating QAC tolerance (Fig. 4). Many of these regulatory systems cross-talk with multiple regulatory systems and may directly or indirectly control additional genes that thwart antibiotic cross-resistance. For example, many significantly altered proteins shared by both BZKR1 and CETR1 were regulated by anaerobic and stress inducible FNR protein (Fig. 4) a transcriptional regulator reportedly controlled by MarA (83). Cooperative actions of CRP-cAMP and FNR have been shown to enhance antibiotic susceptibility in *E. coli* (84). Secondly, there may be a requirement for QAC tolerant species to possess or acquire additional antibiotic resistance genes regulated by stress induction. In our study, *E. coli* K-12 is susceptible to antiseptics as well as antibiotics, but, in past studies of *E. coli* isolates collected from food and clinics these strains may possess additional antibiotic resistance genes that the K-12 strain did not possess. Class I integrons carried on plasmids can transmit QAC selective efflux pump genes in addition to extended spectrum β-lactamase (ESBL) resistance genes and other antibiotic resistant determinants (11, 12, 25, 85); many promoters on these integrons are stress inducible (86, 87). Enterobacterial species exposed to QACs over time may create cells with optimal stress induction systems to not only enhance intrinsic QAC tolerance mechanisms but also create cellular conditions favorable for antibiotic resistance gene co-selection.

## 4.0 Materials and Methods

### 4.1 Chemicals and bacterial strains used in this study

QACs and chemicals used in this study were obtained from Tokyo Chemical Industry America (USA), Millipore Sigma (USA), Fisher Scientific (USA) and VWR (Canada). *E.coli* K-12 BW25113 (88) was obtained from the Coli Genetic Stock Centre (cgsc2.biology.yale.edu). All cultures were grown in Luria-Bertani (LB) broth at 37°C with shaking at 150 revolutions per min (RPM). Cultures grown ‘with selection’ refers to cultures grown at 50% of the final QAC concentration (40 µg/mL BZK; 50 µg/mL CET).

### 4.2 QAC adaptation experiments

QAC adaptation experiments were performed in broth medium using a repeated sub-culturing method similar to Bore *et al*. 2007 (10) with modifications. Briefly, *E. coli* K12 BW25113 was grown overnight (18 hrs) from a dimethylsulfoxide (DMSO) cryopreserved stock and then diluted 10^−2^ into 5 mL LB containing either BZK or CET at 20% of the wildtype (WT) MIC value (3, 6 µg/mL, respectively) in triplicate. The next day, cultures were re-inoculated into 5 mL of fresh LB containing stepwise increases of 2 µg/mL QAC and grown overnight. Cultures with visible growth at the highest QAC concentration were chosen for re-inoculation into culture containing further stepwise QAC increases (2-4 µg/mL steps). This sub-culturing cycle was repeated for a total of 40 days. This process generated six independent replicates: BZKR1, BZKR2, BZKR3; CETR1, CETR2, CETR3 (Table 1). During the adaptation process, a wildtype (WT) culture was also passaged in LB in tandem as a control to determine genetic drift in LB medium after 40 days. At the end of this process, adapted bacteria were cryopreserved in LB with 16% (v/v) glycerol and stored at -80°C.

### 4.3 Antimicrobial Susceptibility Testing (AST) to determine MIC values

A modified broth microdilution AST method (89) was used to determine MIC values. Briefly, cryopreserved stocks of WT or QAC-adapted *E. coli* were inoculated overnight without selection, then overnight with selection. Culture turbidity was measured and standardized spectrophotometrically to obtain an optical density (OD_600nm_) of 1.0 unit in LB with a Multiskan Spectrum microplate reader (Thermo Fisher Scientific, USA). The adjusted cultures were diluted 10^−2^ into 96-well microtiter plates containing 2-fold serial dilutions of antimicrobial in LB broth. AST for all replicates was performed in technical triplicate. A total of 22 antimicrobial compounds were included for AST and are summarized in Table 1. For antimicrobials that required solubilization in ethanol (chlorhexidine; CHX, linezolid; LZD, rifampicin; RMP), methanol (erythromycin; ERY), or DMSO (alexidine; ALX, trimethoprim-sulfamethoxazole; SXT), control plates containing the same concentration of these solvents only were used to assure the results were due to the antimicrobial. Once inoculated, all microplates were incubated overnight before OD_600nm_ spectrophotometric measurement. Significant increases in MIC were defined as >2-fold change as compared to WT.

### 4.4 Growth curve, QAC tolerance stability fitness, biofilm formation assays

Bacterial fitness was assessed for QAC-adapted replicates in 96-well microplate broth cultures. Microplate growth curves were set up as described for AST testing, however, QAC concentrations used in these assays were performed at 10-20% of the WT strain’s MIC value (BZK: 3.6 µg/mL, CET: 3 µg/mL) to compare strains grown at identical drug exposures. Growth curves were measured spectrophotometrically over 24 hrs using a Synergy Neo2 Hybrid Multimode reader (Biotek, USA). 24 hr growth curve experiments were repeated for each replicate tested in a variety of rich (LB, LB + 0.4% (w/v) glucose, cation adjusted Mueller Hinton broth; MHB, Tryptic Soy broth; TSB) and minimal (minimal nine salts; M9, Davis Glucose; DG) media.

The phenotypic stability of each QAC-adapted *E. coli* was assessed by repeated sub-culturing of the QAC-adapted replicates in LB without QAC over 10 days. Each day, AST against the replicate’s respective QAC was performed as described above. All stability experiments were completed in technical triplicate per replicate.

Capacity for biofilm formation was determined using the crystal violet staining protocol of 24 hr MBEC Assay® biofilm devices (Innovotech, Canada) grown in LB without selection outlined in (90). Briefly, each replicate was grown in LB broth overnight, then standardized to an OD_600nm_ of 1.0 in LB and diluted 1:10,000 into the wells of MBEC pegged lid 96-well microtiter plate. Inoculated MBEC plates were incubated for 24 hrs where the pegged lids were rinsed in sterile phosphate buffered saline (PBS) to remove planktonic cells, then stained in 0.1% (w/v) crystal violet (CV) for 5 min. Pegs were de-stained in a fresh 96-well plate containing 200 µl 30% (v/v) acetic acid in sterile water for 10 min then measured at an absorbance of 550 nm (Abs_550nm_) using a Multiskan Spectrum microplate reader (Thermo Fisher Scientific, USA). Each replicate’s biofilm capacity was measured in 7 technical replicates.

### 4.5 Scanning Electron Microscopy (SEM)

To identify morphological anomalies between WT and QAC-adapted *E. coli*, SEM was performed using conditions described by Golding *et al*. 2016 (91). Briefly, bacterial samples were grown with selection to an OD_600nm_ of 0.5 units, pelleted by microcentrifugation (30 sec at 14,000 RPM) and resuspended in PBS. Bacteria were processed and imaged using the ‘gold coated sample preparation’ method described (91) with the sole modification that the bacteria were diluted 1:1000 during the loading of the sample onto the filter. Each sample was imaged in the SEM at 5,000X magnification in five different locations on the filter to help assess overall patterns. Twenty bacterial cell lengths and widths were measured in 5 separate images for each replicate (n=100) measured utilizing ImageJ v1.8.0 (92). Statistical analysis of measured cell lengths and widths were analyzed using Prism6 V6.0 (Graphpad Software, USA) and significant differences were assessed using the Student’s *t*-test (*P* < 0.01).

### 4.6 Whole genome sequencing (WGS), genome assembly, and SNV analysis

Genomic DNA (10-30 ng/µL) was isolated from each QAC-adapted replicate using a Purelink Microbiome DNA isolation kit (A29790, ThermoFisher Scientific, USA) according to manufacturer’s instructions for bacterial culture DNA extraction. Genome sequencing was performed by MicrobesNG (https://microbesng.com/; UK) with an Illumina-MiSeq system (Illumina Inc., USA) at a minimum of 30X coverage. Sequencing details are found in Table S2. Trimmed paired reads were generated and assembled in-house using the MicrobesNG pipeline with *E. coli* BW25113 (CP009273.1) as the mapping reference. SNV analysis was performed using Geneious v11.1.5 software (Biomatters Ltd., New Zealand) to identify SNVs and compare DNA sequences from each QAC-adapted replicate to the reference genome. We controlled for genetic drift of the adapted replicates by eliminating any WT SNVs found in the QAC-adapted replicates. A multiple sequence alignment of genome assemblies was created from the mapped reads, and a Maximum Likelihood tree was constructed using *E. coli* BW25113 (NZ_CP009273) as the root sequence using PhyML v.3.3.20180621 (93). Confidence of tree branching was determined by performing 1000 bootstrap replicates and these values are indicated at each node in the dendrogram.

### 4.7 Proteomic analysis and gene ontology

#### 4.7.1 Sample preparation

For proteomic analysis, BZKR1 and CETR1 replicates were selected for further analysis based on higher total SNVs, mean QAC MIC values, and growth characteristics when compared to the WT and other adapted strains. All cultures were grown in 4 L batches with selection to a final OD_600nm_ = 0.5 units. Samples were centrifuged at 6000 RPM for 10 mins (Avanti-J-E HPC, Beckman, USA), and the pellet was resuspended in an equal volume of isolation buffer (50 mM 3-(N-morpholino) propanesulfonic acid (MOPS), 8% v/v glycerol, 5 mM ethylenediaminetetraacetic acid (EDTA), 1 mM dithiothreitol, pH 7). Cell pellets were homogenized in sterile Milli-Q water, mixed with 100 µl 0.1-mm glass beads (Scientific Industries Inc., USA), and heated for 5 min at 95°C. Cells were lysed via highspeed vortex-mixed for 3 min, followed by 1 min centrifugation at 3000 RPM centrifugation. The supernatant was collected and cold sterile Milli-Q water was added to the beads, followed by 1 min vortex/ 1 min centrifugation, where the protein supernatant was pooled. This was repeated 5 times to extract the remaining protein, which was stored at -80°C. Protein was quantified using a bicinchoninic acid (BCA) protein assay kit, with bovine serum albumin (BSA) as the standard (Pierce Protein Research Products; Thermo Fisher Scientific, USA). 100 µg of protein was isolated from BZKR1 and CETR1 in biological triplicate, and digested with trypsin (Promega, USA) overnight (16-18 hrs) using a filter-assisted sample preparation (FASP) method described previously (94). Following digestion, all samples were dried down and reconstituted using mass spectrometry grade water to a final concentration of 1 µg/µl for LC-MS/MS analysis.

#### 4.7.1 Nanoflow-LC-MS/MS

Each sample was separately analysed using a nano-flow Easy nLC 1200 connected in-line to an Orbitrap Fusion Lumos mass spectrometer with a nanoelectrospray ion source at 2.3 kV (Thermo Fisher Scientific, USA). The peptide samples were loaded (2 µl) onto a C_18_-reversed phase Acclaim PepMap 100 trap column (2 cm x 75 µm, 3 µm particles; Thermo Fisher Scientific, USA) with 30 µL of buffer A (2% v/v acetonitrile, 0.1% v/v formic acid) and then separated on an Easy Spray column (50 cm long, 75 µm inner diameter, 2 µm particles; Thermo Fisher Scientific, USA). Peptides were eluted using a gradient of 2-30% buffer B (80% v/v acetonitrile, 0.1% v/v formic acid) over 100 min, 30%-40% buffer B for 20 min, 40%-100% buffer B for 5 min and a wash at 100% B for 10 min at a constant flow rate of 250 nl/min. Total LC-MS/MS run-time was about 175 min, including the loading, linear gradient, column wash, and the equilibration.

LC-MS/MS data was acquired using the settings described below. The top most abundant precursor ions from each survey scan that could be fragmented in 1 second (s) were dynamically chosen, where each ion was isolated in the quadrupole (0.7 *m/z* isolation width) and fragmented by higher-energy collisional dissociation (27% normalized collision energy). The survey scans were acquired in the Orbitrap at mass over charge ratios (*m/z*) of 375-1500 with a target resolution of 240,000 at *m/z* 200, and the subsequent fragment ion scans were acquired in the iontrap at a rapid scan rate. The lower threshold for selecting a precursor ion for fragmentation was 1.5 × 10^4^. Dynamic exclusion was enabled using a *m/z* tolerance of 10 parts per million (ppm), a repeat count of 1, and an exclusion duration of 15 s.

#### 4.7.1 Data processing

All spectra were processed using MaxQuant (v1.6.7, Max Planck Institute) using the imbedded Andromeda search engine. Searches were performed against a subset of the SwissProt database set to *E. coli* K-12 (4519 sequences). The following search parameters were used: Carbamidomethyl (C) was selected as a fixed modification, Oxidation (M) and Acetyl (Protein N-term) as variable modifications, fragment ion mass tolerance of 0.5 Da, parent ion tolerance of 20 ppm, and trypsin enzyme with up to 2 missed cleavage. False discovery rates were set up using 0.01 for peptides, 0.01 for proteins, and at least 1 razor peptide per protein. Label free quantification (LFQ) was enabled for Quantitation. Resulting LFQ intensities were imported into Perseus v1.6.5 (Max Planck Institute)(95). In Perseus the data was Log2 transformed. Then all the proteins that did not have a least 3 valid log2 LFQ intensities from ID were filtered out. Proteins present were assessed for significance using volcano plots with a modified Student’s t-test (false discovery rate of 0.05; S_0_= 0.1). Enrichment analysis of protein to protein interactions was performed with Cytoscape v3.7.2 (96) using the StringApp v1.5.0 software package (97). To analyze functional pathways involved in adaptation, significant proteins were categorized using the KEGG pathway database with ClueGO v2.5.5 (98) application for Cytoscape. Default parameters for KEGG analysis were used, with network specificity at “medium” and 50% overlap of genes for the group merge.

## Acknowledgements

We would like to thank Stuart McCorrister and the Mass Spectrometry and Proteomics Core Facility group (National Microbiology Laboratory, Public Health Agency of Canada, Winnipeg, Manitoba, Canada) for help with proteomics data acquisition and analysis. Thanks to the Diagnostic Microscopy and Imaging group (National Microbiology Laboratory, Public Health Agency of Canada, Winnipeg, Manitoba, Canada) for assistance with scanning electron microscopy experiments. Funding for this study was provided by Natural Sciences and Engineering Research Council of Canada (NSERC) Discovery Grant (DG) [RGPIN-2016-05891] operating grant to DCB.

## Author Contributions

DCB and NHC designed the study and NHC performed the adaptation experiment. NHC and KACG gathered the MIC data. KACG and BSJG gathered growth curve data. KACG gathered the stability data. Genomic DNA extractions and SNV identification were performed by NHC, and SLR generated and analyzed the phylogenetic trees. BSJG and SLR completed the SEM imaging with DRB, SLH, and TFB. BSJG and SLR prepared proteomic sample preparations for LC MS/MS collection by PMC and GRW. CJS performed biofilm assays. KACG, BSJG, and SLR analysed all data and prepared manuscript figures. KACG, BSJG, and SLR wrote the manuscript drafts in consultation with DCB, where DCB and GGZ edited. All authors read and approved the final manuscript.

## Figure Captions

**Supplementary files**. Figures S1 to S4 and Tables S1-S3 are provided as attached supplementary data files.

